# The 8-nm spaghetti: well-structured glycans coating linear tetrapeptide repeats discovered from freshwater with CryoSeek

**DOI:** 10.1101/2024.12.15.627649

**Authors:** Tongtong Wang, Yitong Sun, Zhangqiang Li, Nieng Yan

## Abstract

We recently developed a research strategy, termed CryoSeek, to identify uncharacterized bio-entities from natural or endogenous resources using cryo-electron microscopy (cryo-EM). Here we report the discovery of a glycofibril whose primary molecular mass is attributed to a thick glycan shell. The 3.3-Å resolution cryo-EM reconstruction reveals that the only protein component of the glycofibril, which is approximately 8 nm in diameter, is a linear chain of tetrapeptide repeats. Each tetrapeptide repeat consists of a 3,4-dihydroxyproline (diHyp), a Ser or Thr, and two less conserved residues. Two and one glycan chains are respectively O-linked to the diHyp and Ser/Thr residues. The protein sequence pattern of this glycofibril is similar to that of our recently observed TLP-4, although the glycan chains are different. We rename the previously characterized glycofibril as TLP-4a and designate this one as TLP-4b. Our discoveries reveal the critical role of glycans in structural folding of glycoconjugates and shed light on understanding the carbon/nitrogen ratio in biospheres.

## Introduction

Carbohydrates, the most abundant organic molecules on Earth, are one of the four fundamental biomolecules essential for all organisms (1, 2). They function in a wide range of biological activities, such as serving as the primary energy source for the majority of species, participating in the construction of cells, transducing signals, and modulating the folding and functions of proteins and nucleotides (1-8). Recent technological advancements in structural biology have significantly expanded our understanding of proteins (9-14). However, structural biology has yet to provide any major insights into the biological characterization of carbohydrates due to the lack of high-resolution glycan structures. The technical challenges in structural biology of glycans come from the compositional diversity of monosaccharides, branching complexity, and conformational flexibility of polysaccharides (15, 16).

Earlier this year, we have introduced a research strategy, named CryoSeek, which employs cryo-electron microscopy (cryo-EM) to search for unknown bio-entities from any native or endogenous samples (17). As a proof of concept, we identified two protein fibrils in the water sample from the Tsinghua Lotus Pond that are likely to be surface pili from uncharacterized bacteria (17). Most recently, we identified an unprecedented glycofibril, in which the only protein portion is a linear peptide chain and the surrounding glycan shell constitutes more than 90% of the molecular mass. The assembly of the fibril is completely supported by the packing among glycans. We named this fibril TLP-4 for its isolation from the <u>T</u>singhua <u>L</u>otus <u>P</u>ond and the central stem being made of <u>tetra</u>peptide repeats (18). Among the four amino acid residues in each repeat, two consecutive ones are conserved, a 3,4-dihydroxyproline (DiHyp) with 3-OH and 4-OH being O-glycosylated and an adjacent serine/threonine (Ser/Thr) with a third large O-linked glycan. The other two residues are less conserved.

Through bioinformatic analysis, we found a large number of protein sequences containing such tetrapeptide repeats in the NCBI Non-redundant (NR) database, suggesting that TLP-4 like glycofibrils might exist widely across species (18). While we were setting up lab culture systems for various diatom species to systematically characterize TLP-4, another glycofibril isolated from the pond water with the same sequence pattern as TLP-4 was structurally resolved. Despite identical PS/Txx repeats with three glycan chains linked to DiHyp and Ser/Thr, the number of sugar residues in this glycofibril is substantially larger than that in TLP-4, with distinct branches. To distinguish these two types of glycofibrils, we will refer to the previous and current ones as TLP-4a and TLP-4b.

As TLP-4b was characterized after the acceptance of the manuscript on TLP-4a, we hereby prepare this preprint to complement the study that is in press. Both TLP-4a and TLP-4b are approximately 8 nm in diameter and composed mainly of carbohydrates. In a sense, they are reminiscent of rice noodles or spaghetti in the microscopic world.

## Results

### Discovery of TLP-4b from the Tsinghua Lotus Pond

Following the same experimental procedures for CryoSeek as previously described (17, 18), we successfully obtained the 3D reconstruction of TLP-4b, with an overall resolution of 3.3 Å. Although TLP-4b was assigned to different groups during 2D class averaging and 3D reconstruction, its helical parameters are nearly identical to those of TLP-4a. The helical rise between the adjacent asymmetric units is 12.4 Å for both glycofibrils, and the twist is 134.0° for TLP-4a and 138.6° for TLP-4b (Fig. 1A).

**Figure 1.**
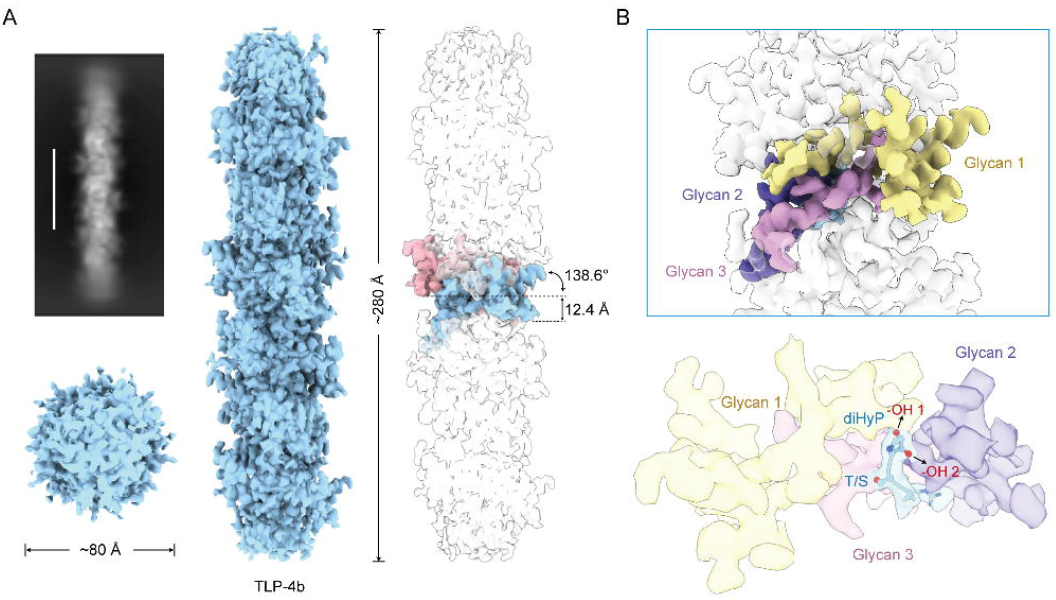
TLP-4b consists of linear tetrapeptide repeats coated with dense glycans. (*A*) High-resolution 3D reconstruction of TLP-4b with helical parameters. *Left*: Two perpendicular views of segmented TLP-4a and the corresponding 2D average. *Right*: Two adjacent asymmetric units. The two units are colored pink and light blue. The helical rise and twist for TLP-4b are 12.4 Å and 138.6°, respectively. The overall resolution is 3.3 Å. The EM map is contoured at 4.5 σ. (*B*) Three glycan branches of TLP-4b. *Upper*: Each helical repeat of TLP-4b has three branches of glycan densities linked to the central peptide. *Lower*: Glycan 1 and 2 are O-linked to a 3,4-dihydroxyproline (diHyp) residue, while the neighboring glycosylated residue can be a Thr/Ser. The EM map is contoured at 9 σ. All structural figures were prepared in ChimeraX (31).

After scrutinizing the 3D EM map of TLP-4a for nearly one year, we could easily recognize the glycan-linked 3,4-dihydroxyproline (diHyp) and the adjacent Ser/Thr in each tetrapeptide repeat of TLP-4b. The other two residues are still less conserved, resulting in poor resolution for the local map. Two glycan chains, hereafter referred to as glycan 1 and glycan 2, are bonded to the 4-OH and 3-OH groups of diHyp, respectively, while the third glycan chain is O-linked to Ser/Thr (Fig. 1B).

### TLP-4b contains more sugar residues in each glycan chain

The identities of these three glycan chains are yet to be accurately assigned at the present resolution. Therefore, we tentatively modeled glycosyl moieties to indicate the glycan branches and orientations (Fig. 2A). Similar to TLP-4a, the local resolutions drop from the center to the periphery. We only attempted to model in sugar residues to densities where each individual moiety could be distinguished. Therefore, the actual number of sugar residues should exceed what can be structurally modeled.

**Figure 2.**
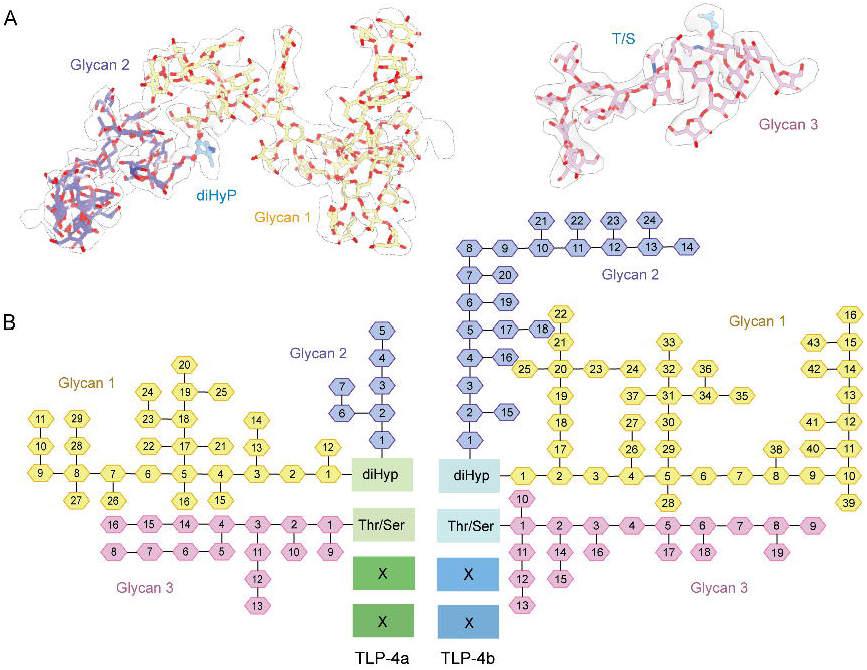
Modeling of glycans in each tetrapeptide repeat of TLP-4b. (*A*) Representative EM densities of the three glycan chains in each tetrapeptide repeat. A total of 86 glycosyl moieties have been modeled in each repeat of TLP-4b. Glycans 1, 2, and 3, colored yellow, blue, and pink, are assigned 43, 24, and 19 sugar residues, respectively. (*B*) Comparison of the glycan chains in each repeat of TLP-4a and TLP-4b. The hexagons represent unknown glycosyl moieties. The number and branching patterns for the three glycans in each repeat vary between TLP-4a and TLP-4b. The schematic illustration here represents only the number and branching patterns of glycans in TLP-4a and TLP-4b, not their chirality.

In total, we were able to assign 86 glycosyl moieties in each repeat, with 43, 24 and 19 sugar residues in glycans 1, 2, and 3, respectively. In comparison, we were only able to build 52 glycosyl moieties for each repeat of TLP-4a, with 29, 7, and 16 sugar residues in the three glycans (Fig. 2).

### Glycans are responsible for structural “folding”

As in TLP-4a, the overall appearance of the glycofibril is determined by the glycans. Water molecules, which are important for mediating indirect hydrogen bonds among sugar residues, are invisible in the 3.3 Å resolution map. In addition, the lack of accurate glycosyl assignment prevents a reliable analysis of the sugar interactions. Nevertheless, the structure affords a glimpse of the “folding” of the glycan branches (Fig. 3).

**Figure 3.**
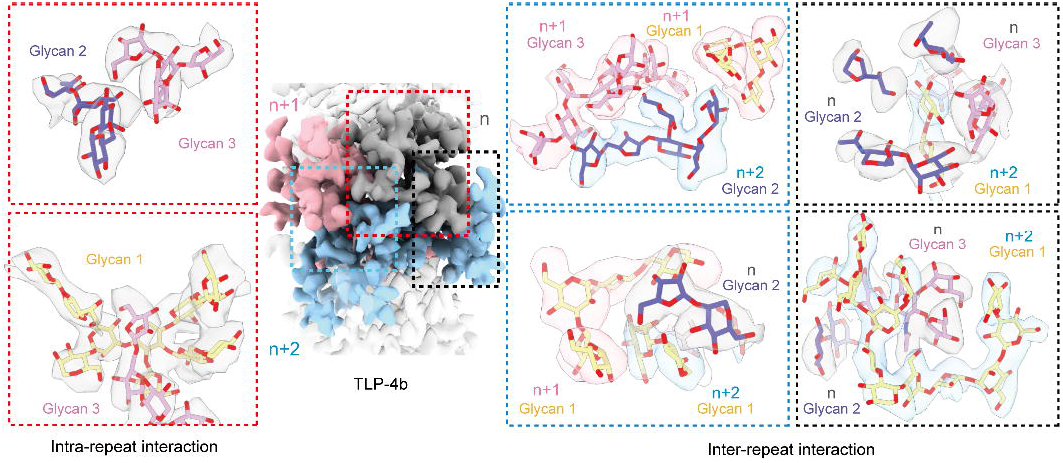
Glycans are responsible for structural assembly of TLP-4b. EM densities for three adjacent repeats are shown and labeled n (grey), n+1 (pink), and n+2 (ligh blue). The extensive intra- and inter-repeat interactions mediated by glycans are illustrated in the *left* and *right* insets. The EM map is contoured at 9 σ.

The glycans intertwine to support a stable structural assembly of the fibril. The three glycan chains in each tetrapeptide repeat of TLP-4b form multiple interactions with one another. The intra-repeat sugar interactions primarily involve direct or water-mediated hydrogen bonds and stacking of the sugar rings (Fig. 3, left two panels). The inter-repeat interactions are even more compact and extensive, with four representative interfaces (Fig. 3, right four panels). These extensive glycan interactions collectively determine the structure of the well-ordered helical fibril of TLP-4b.

## Discussion

In this study, we determined the structure of another heavily glycosylated fibril from water samples collected from the Tsinghua Lotus Pond. Despite significant differences in the number and branching patterns of the sugar moieties, it is the glycans that determine the fibril assembly for both TLP-4a and TLP-4b. The identification of TLP-4b supports our previous bioinformatic analysis that suggested widespread existence of similar glycofibrils across organisms (18).

DiHyp, primarily identified in diatoms (19), serves as a center for simultaneously anchoring two bulky glycan chains. The next residue is Ser or Thr, either of which is heavily glycosylated. The three glycans chains in every four amino acids form a highly ordered and tightly packed glycoshell that seamlessly protects the polypeptide core from proteases, allowing TLP-4 fibrils to remain stable in the natural environment.

As for the potential origins and functions of TLP-4a/b, some clues come from the characterizations of the extracellular polymeric substances (EPS), which are secreted by certain benthic diatoms (20-22). They are reported to be particularly rich in PTSA/G repeats and acidic polysaccharides (22), suggesting a possible origin for TLP-4a/b. The functions of EPS components are diverse, including adhesion, biofilms cohesion, water retention, and absorption of polar organic compounds. They can serve as nutrient sources, and act as a sink for excess energy (23).

Despite these clues, the most intriguing question accompanying the discovery of the TLP-4 glycofibrils concerns the unknown enzymes, as well as potential chaperons, that catalyze the synthesis and assembly of the highly dense glycans around the linear tetrapeptide repeats. In addition to the number and types of glycosyltransferases, it is completely unclear how and where these glycosylation modifications occur. Of note, TLP-4a and TLP-4b, which share a similar protein core sequence, have different glycan compositions and structures. The determinants for these differences remain to be discovered.

As these fibrils were isolated from a natural environment and the only sequence information comes from the degenerate tetrapeptide repeats, it is literally impossible to identify the enzymes using any conventional approach. Model organisms need to be established for genetic manipulations. Data mining of the experimental and predicted structures may also provide important clues (24-26). Identification of the related enzymes and chaperons may facilitate materials science, synthetic biology and hybrid synthesis of glycan foldamers that could have various applications.

It is noted that the peripheral densities in TLP-4a and TLP-4b were not structurally assigned with sugar moieties owing to the low resolution. Therefore, the actual mass ratio of the sugar and amino acid residues exceeds 95:5 in both TLP-4 fibrils. Such glycofibrils may afford a means for carbohydrate storage in addition to starch, glycogen, and cellulose (27, 28). Unlike starch and glycogen, which are localized within the cell, or cellulose, which define the cell boundary, the glycofibrils are outside the cell body and able to extend into unlimited extracellular space, offering an important route for storing the fixed carbon. These fibrils may serve as nutrients for the organisms that produce them or other species. On the other hand, the distinct ratio of sugar residues and amino acid residues in these glycofibrils may provide a regulatory mechanism to balance the fixation of carbon and nitrogen.

In sum, our studies presented here, along with that on TLP-1 and TLP-4a, exemplify the power of CryoSeek in discovering completely uncharacterized bio-entities, which serve as the starting point for a series of hypothesis-driven studies.

## Materials and Methods

### Data processing

The pre-treatment of water from the Tsinghua Lotus Pond, cryo-EM sample preparation, and data acquisition procedures were nearly identical to those previously reported (17, 18). Regarding the data processing of TLP-4b, in brief, a total of 77,044 good particles were selected after deep 2D classification. Following ab-initio 3D map reconstruction and refinement, the resulting map was used for helical symmetry search. The helical rise and twist of TLP-4b were initially identified as 12.34 Å and 138.70°, respectively. After heterogenous refinement, a total of 44,738 particles remained, yielding a map with a global resolution at 3.29 Å, and the helical rise and twist determined to be 12.37 Å and 138.63°, respectively.

The reported resolution for TLP-4b was calculated based on the gold-standard Fourier shell correlation (FSC) 0.143 criterion using cryoSPARC (29, 30).

### Model building and refinement

Model building of TLP-4b was carried out based on the 3.3 Å reconstruction maps. Since the protein core of TLP-4b is similar to that of TLP-4a, the atomic coordinates of TLP-4a were fitted into the EM map of TLP-4b in ChimearX (31). We constructed hexoses or pentoses based on the EM map as the actual identities of these glycans could not be determined. Finally, we manually built the atomic model of TLP-4b in COOT (32).

The final model of TLP-4b was refined using PHENIX with secondary structure and geometry restraints in real space (33). The structure was validated through examination of the Clash scores, MolProbity scores, and statistics of the Ramachandran plots in PHENIX (33, 34).

## Data and code availability

The cryo-EM map of TLP-4b has been deposited in the Electron Microscopy Data Bank (https://www.ebi.ac.uk/pdbe/emdb/), under the accession codes EMD-xxxxx. The corresponding atomic coordinate has been deposited in the Protein Data Bank (http://www.rcsb.org), under the accession code xxxx.

## Author contributions

N.Y. and Z.L. conceived the project. T.W., Z.L., and Y.S. performed experiments and collected cryo-EM data. T.W., Z.L., and Y.S. processed the cryo-EM data and determined the structure. T.W., Z.L., Y.S., and N.Y. did structural analysis. N.Y., Z.L. and T.W. wrote the paper.

## Acknowledgements

We thank Dr. Xiaomin Li and Dr. Jianlin Lei for technical support during EM image acquisition. We thank the Tsinghua University Branch of China National Center for Protein Sciences (Beijing) for providing the proteomics facility and cryo-EM facility support. We thank the computational facility support on the cluster of Bio-Computing Platform (Tsinghua University Branch of China National Center for Protein Sciences Beijing). This work was funded by the National Natural Science Foundation of China (project 92478205 and 32330052 to N.Y.).

